# Dual GLP-1/FGF21 agonism suppresses voluntary alcohol consumption, alcohol choice, and nucleus accumbens dopamine modulation

**DOI:** 10.64898/2026.04.30.721773

**Authors:** Bart J. Cooley, Parul Sirohi, Caslin A. Gilroy, Joy (Shijia) Tong, Catherine G. Price, Emily Mitchell, Willow A. Heller, Ashutosh Chilkoti, Andrew J. Lawrence, Gavan P. McNally, E. Zayra Millan

**Affiliations:** School of Psychology, UNSW Sydney, Australia; Department of Biomedical Engineering, Duke University, Durham, NC 27708, USA; The Florey Institute of Neuroscience and Mental Health, University of Melbourne, Parkville VIC, Australia

## Abstract

Excessive alcohol consumption remains a major public health challenge with limited therapeutic options. Both glucagon-like peptide-1 (GLP-1) and fibroblast growth factor-21 (FGF21) independently regulate alcohol intake through complementary metabolic and reward pathways, but their combined potential has not been explored. Here, we report that a long-acting dual agonist, GLP1-ELP-FGF21 modulates behavioural, neurophysiological, and cognitive components of alcohol seeking in mice. A single GLP1-ELP-FGF21 dose reversibly reduces voluntary alcohol intake for at least 72 hours in male mice, has sustained effects in female mice, and markedly blunts nucleus accumbens dopamine transients aligned to the initiation and termination of lick bouts during alcohol consumption. To assess its effects on decision-making, we used a novel two-choice (alcohol versus food) decision task modelled with evidence-accumulation frameworks. Alcohol choice behaviour conformed to evidence accumulation decision models: Linear Ballistic Accumulator (LBM) and Racing diffusion models (RDM). Critically, GLP1-ELP-FGF21 selectively reduces choices for alcohol and slows the latent accumulation rate for alcohol options, without affecting food-directed choice or non-decision processes. Sensory-specific satiety devaluation confirms that reductions in reward value are explained by reductions in accumulation rates. Together, these results highlight GLP1-ELP-FGF21 as a therapeutic strategy for alcohol use disorder via modulation of central reward pathways and decision-making when confronted with alcohol reward

## Introduction

Excessive alcohol consumption is a major contributor to preventable morbidity and mortality worldwide, driving liver disease, cardiovascular pathology, cancer, and neuropsychiatric disorders. Despite its substantial public health impact, pharmacological treatments for alcohol use disorder (AUD) remain limited by modest efficacy and poor tolerability. Approved therapies—including naltrexone, acamprosate, and disulfiram—produce only small effect sizes and suffer from poor adherence, underscoring the need for mechanism-based interventions that directly target the neurobiological drivers of alcohol consumption and reward (Heilig et al., 2024; Jonas et al., 2014).

Incretin and neuroendocrine-based therapies have recently emerged as promising candidates (Brockway & Crowley, 2024; Heilig et al., 2024). Among these, glucagon-like peptide-1 (GLP-1) receptor agonists have generated considerable interest. Preclinical studies show that GLP-1 agonists reduce alcohol-seeking and intake (Cajsa Aranäs et al., 2023; Díaz-Megido & Thomsen, 2023; Egecioglu et al., 2013), and early clinical studies involving GLP-1 mimetics such as semaglutide and liraglutide, report reduced alcohol craving and consumption—sometimes at doses lower than those used for obesity—with effect sizes that compare favourably to existing treatments (Hendershot et al., 2025; Klausen et al., 2022). These behavioural effects align with mechanistic evidence showing that GLP-1 acts on key mesolimbic and thalamic pathways associated with reward and alcohol motivated behaviour (Chuong et al., 2023; Dixon et al., 2020; Klausen et al., 2022; Ong et al., 2017; Subhani et al., 2024).

Efforts to enhance GLP-1 efficacy have increasingly focused on combination strategies involving other peptide agonists (Gilroy et al., 2020; Oertel et al., 2024; Pan et al., 2021; Rosenstock et al., 2021) or small molecules (Petersen et al., 2024), although their effect on alcohol consumption remain largely unexplored. Among potential co-targets, fibroblast growth factor-21 (FGF21), is particularly compelling. Alcohol intake elevates circulating FGF21 in humans (Lanng et al., 2019) and single nucleotide polymorphisms (SNPs) in FGF21 or its obligate co-receptor β-Klotho (KLB) are associated with AUD risk (Schumann et al., 2016). Preclinical studies further show that FGF21 suppresses alcohol intake: FGF21 overexpression reduces alcohol preference (Talukdar et al., 2016), whereas KLB deletion increases consumption (Schumann et al., 2016), and pharmacological administration of FGF21 suppresses alcohol consumption in mice and non-human primates (Flippo et al., 2022). FGF21 may also confer hepatoprotective benefits relevant to clinical AUD populations (Harrison et al., 2024; Nielsen et al., 2023).

Motivated by recent advances in GLP1–FGF21 fusion agonists (Gilroy et al., 2020; Pan et al., 2021), we evaluated a unimolecular GLP1–FGF21 agonist (Gilroy et al., 2020) in which the two peptides are fused via an intrinsically disordered elastin-like polypeptide (ELP) that enables sustained zero-order drug release of the dug over a week (Gilroy et al., 2020). We first assessed its effects on alcohol consumption and on dopamine signalling in the nucleus accumbens shell (AcbSh) during alcohol intake. To strengthen translational relevance, we then tested this compound in a discrete choice alcohol task and applied evidence-accumulation models commonly used in human decision-making research. This cross-species computational approach enables identification of conserved cognitive mechanisms through which the dual-agonist may alter alcohol-motivated behaviour.

## Materials and Methods

### Subjects

Adult C57BL/6J mice (OzGene, Perth, Australia) were used. During 24 hr homecage alcohol access, mice were single-housed on a 12:12-hour light/dark cycle (lights on at 07:00 for all experimental studies) and had *ad libitum* access to food and water. For the fiber photometry study, mice were water-restricted (∼18hr) once per week prior to each test. There were no other dietary or fluid restrictions in place. For the alcohol choice study, mice were food restricted and maintained at approx. 85-90% free feeding body weight during instrumental behavioural testing. Behavioural testing occurred during the light phase. All procedures were approved by the Animal Care and Ethics Committee at UNSW Sydney and all attempts were made to minimise the number of animals used.

### Surgery

To monitor *in vivo* AcbSh dopaminergic activity, mice underwent stereotaxic surgery for delivery of the genetically encoded dopamine sensor, GRAB-DA3, and implantation of an optic cannula fibre. Mice were deeply anaesthetised with 5% v/v isoflurane in oxygen-enriched air, secured into a stereotaxic frame (Model 1900, Kopf Instruments) and maintained on 1%-2.5% v/v isoflurane for the duration of surgery. Carprofen (5mg/kg, s.c., Rimadyl, Zoetis) was administered for analgesia, Marcaine (incision site, 0.1 ml (0.5%), s.c.) for local anaesthesia, and ophthalmic gel (Viscotears, Alcon) was applied to avoid corneal drying. The dopamine sensor (pAAV-hSyn-GRAB-gDA3m, Addgene) was then delivered into the right or left AcbSh (+1.4mm A-P; ±0.5 M-L, -4.1 to -4.5 D-V) using a 33-gauge Hamilton syringe controlled via a microinjection pump (World Precision Instruments; 0.5ul total volume; 0.1ul/min rate of delivery). A 400-μm multi-modal optic fibre encased within a ceramic ferrule was implanted above the virus injection site and secured to the skull with stainless-steel screws and dental adhesive. Following surgery, mice received the antibiotic Duplocillin (0.15 ml/kg, s.c.) and were closely monitored until full recovery. All coordinates are reported relative to bregma using the Paxinos and Franklin’s Mouse Brain in Stereotaxic Coordinates [28].

### Homecage drinking apparatus

Mice consumed alcohol in the homecage from sippers constructed from 15ml centrifuge tubes fitted with Hydropac valves (Allied Scientific Products, Australia), sealed with rubber O rings and suspended to one side wall of the homecage via a 3D-printed holder. Two sippers were simultaneously mounted on each drinking day - one contained alcohol solution, the other water.

### Lickometer apparatus

To monitor individual licks during alcohol drinking we used a Med Associates contact lickometer housed in a wide Med Associate chamber (21.59 ([length] x 18.08 [width] x 12.7 [height] cm) with clear Perspex walls and stainless-steel grid floors. The chamber was enclosed in a light and sound attenuating cabinet. Mice accessed alcohol solution (15% v/v) via a stainless-steel spout protruding from the center of the right-side wall of the chamber. Licks were detected via closed-loop contact between the spout and grid floor (Med Associates ENV-250 Contact Lickometer Controller). All experimental events were controlled by Med Associates software.

### Choice apparatus

Operant behavioral training and testing were conducted in standard Med Associates chambers with clear Perspex walls and stainless-steel grid floors enclosed in light and sound attenuating cabinets. Each box contained two retractable levers each side of a central recessed port containing a separated food and liquid trough. 15% v/v alcohol was delivered into the magazine via silicone tubing fitted to a 60mL syringe on a syringe pump. Grain pellets (20mg, Bio-serv #FO163) were delivered by a pellet dispenser. Magazine entries were detected by an infrared photobeam.

### Fiber photometry recording

To measure *in vivo* recording of DA transients during alcohol drinking, mice were tethered to an optical patch cable (RWD) connected to a fiber photometry recording system (RWD R810 Dual Colour Multi-Channel Fiber Photometry System, RWD Life Science Co., Ltd., Guangdong, China). Two alternating excitation wavelengths, 470 nm and 410 nm (isosbestic control signal) were emitted from an LED light source, channelled via the patch cable into a multi-mode optic fiber cannula implant on the mouse’s head. Emitted fluorescence was detected via a sCMOS camera housed within the fiber photometry recording system. Lick event timestamps were synchronized to the recordings via lick-paired transistor-transistor logic (TTL) input signals from Med Associates to the fiber photometry recording system. LED illumination was triggered at the start of each session, controlled similarly by Med Associates. GRAB-DA3 fluorescence was recorded under the blue (470nm) light and control isosbestic signals were detected using violet (410nm) light. Recording parameters were held constant across animals using the RWD software (frame rate = 100 fps; exposure = 7.37 ms; gain = 1; power = 7-10%; LED ∼ 40µW at tip of patch cable).

### Drugs

Drinking alcohol was prepared using 100% ethanol (ChemSupply, Australia) and diluted to 15% v/v with tap water. GLP1-ELP-FGF21 was prepared as described in Gilroy et al. (2020). Exenatide was prepared from frozen stock solutions using sterile saline. We used a 2.5μg/kg dose, selected on the basis of prior work with alcohol drinking mice (Egecioglu et al., 2013). Vehicle was sterile saline (0.9% v/v).

## Experimental Procedures

### Homecage intermittent access to alcohol

Mice had access to two bottles (15% v/v alcohol and tapwater) on an intermittent schedule (3 days/week) (Millan et al., 2017). One bottle contained 15% v/v alcohol and the other tap water. Bottle positions were alternated to prevent side preference. Intake was measured by weighing bottles before and after each session. Mice with low levels of alcohol intake (average intake <1g/kg) were excluded from further testing (Experiment 1: n = 1; Experiment 2: n = 2). Data were excluded if bottles leaked (Experiment 3 water intake at 24 hrs: n = 2; at 72 hrs: n = 1; at 144 hrs: n = 3).

### Lickometer access to alcohol

After three weeks of homecage intermittent access to alcohol mice commenced brief lickometer access to alcohol (1 x 25 min session/week) in addition to their scheduled homecage intermittent access to alcohol (3 days/week). During lickometer sessions, mice were tethered to an optical patch cable and placed in the apparatus chamber for 25 minutes. The first five minutes served as a habituation period. Subsequently, a metal panel was lifted to reveal a single alcohol sipper from which mice were free to drink. Homecage intermittent access to alcohol occurred on the evening following each lickometer session. Drug testing occurred from Week 4 onwards.

### Alcohol choice

Mice first received two weeks of homecage intermittent access to alcohol followed by two weeks of restricted 2 hour intermittent access to habituate them to alcohol during the light-phase. Mice were then food restricted, receiving 1 hour of daily food allotment. They first received a single session of two bottle alcohol access under postprandial thirst conditions. Specifically, animals were provided with their 1 hour food allotment with water removed. After 1 hour the food was removed and animals were provided with two bottles, one containing alcohol and the other water. Food restriction was maintained throughout the experiment and all subsequent instrumental training and choice testing occurred under postprandial conditions on an intermittent schedule.

### Instrumental training

Mice received 2 sessions of magazine training to 15% v/v alcohol and grain which was delivered on a random interval (RI) schedule (RI30 for each reward). Instrumental training involved 8 x 60 min sessions of training to respond for alcohol and grain rewards in blocks where each lever was extended for 15 minutes (maximum of 10 rewards per block). Lever side-reward pairings were counterbalanced across mice. During these sessions mice were phased through an increasing random ratio (RR) criteria beginning with continuous reinforcement (CRF), then RR2, RR5 and RR10 (minimum of two RR5 and RR10 sessions). Mice progressed to the next schedule once they received all 10 rewards in a single block for each reward.

### Free-operant choice

Once mice acquired stable pressing on both levers at RR10 they received 2 x free-operant choice tests with both levers extended. During the first test only, no rewards were delivered in the first 5 minutes of the first test, which served as a non-reinforced probe. For the remaining free-operant choice tests, lever choice was reinforced under RR10.

### Discrete choice training

Mice were trained for 3 sessions on a discrete choice test between alcohol and grain rewards (Augier et al., 2018). This involved a task structure with two successive phases: a reward sampling phase involving 4 sampling trials (2 x trials per lever) and a choice phase (40 trials). During the sampling phase, one lever was extended at a time with the alcohol lever always presented first. Discrete choice trials involved the concurrent presentation of both levers. A lever press resulted in the delivery of the paired reward and retraction of both levers for 1 minute after a choice was made. Animals were allowed 20 seconds to make a choice, otherwise the trial was considered an omission and both levers were retracted.

### Discrete choice test with treatment

Animals were excluded from test due to technical errors or low preference (n = 6). Mice were treated with GLP1-ELP-FGF21 (1000 nmol/kg) or vehicle. They then received 2 discrete choice tests.

### Devaluation

To assess the goal-directed nature of the choice behaviour, outcomes were devalued using a within-subject specific-satiety manipulation. Half of the mice (n = 4) received one hour of food allotment before gaining 20-minute access to 15% v/v alcohol in the home cage with chow removed. The other half of the animals received one hour of food allotment with grain pellets for one hour and 20 minutes.

### Drug Testing Procedure

For homecage alcohol drinking tests, mice received subcutaneous injections approx. 1 hour prior to placement of the sipper apparatus into the homecage. For lickometer access tests, injections occurred approx. 1 to 3 hours prior to placement in the lickometer chamber. For alcohol choice tests, injections were administered approx. 3 to 4 hours prior to placement in the self-administration chamber. For tests involving multiple treatments or doses, treatment conditions were allocated in a repeated measures latin-square design.

### Histology

Mice were anesthetised with intraperitoneal injections of sodium pentobarbital (100 mg/kg) and perfused with 0.9% saline solution containing 1% sodium nitrate and heparin (5000 IU/ml), followed by phosphate buffer (PB) solution (0.1 M) with 4% w/v PFA. Brains were extracted, incubated in 20% w/v sucrose solution in PB for cryoprotection, sliced coronally (40μm) using a cryostat, and stored in PB solution with 0.1% sodium azide at 4°C. Optic fiber placements and virus expression were determined via immunohistochemistry. Sections were washed in PB with 0.9% saline (PBS; 0.1 M, pH 7.2, 3 x 10 min washes), incubated in buffered blocking solution (0.5% Triton-X, 10% normal goat serum (NGS) in PBS) for 1hr, then incubated overnight in chicken antiserum against GFP (1:2000 in 0.2% Triton-X, 2% NGS in PBS; Invitrogen). Tissue was then washed with PBS (3 x 10 min washes), incubated for 2hrs in goat anti-chicken secondary antibody, AlexaFluor-488 (1:1000 in 0.2% Triton-X, 2% NGS in PBS; Abcam, catalog #ab150169), washed again with PBS (3 x 10 min washes), then stored in PB solution with 0.1% sodium azide at 4°C prior to mounting onto gelatinized slides. Slides were left to dry, cover-slipped, then imaged using a slide scanner (Axioscan 7, ZEISS).

## Data Analysis

Group numbers are indicated in Results. All analyses were conducted using MATLAB, R (R Core Team, 2025), SPSS, Psy statistical packages. Inferential statistics were based on a planned set of orthogonal repeated measures comparisons with statistical significance set at *p* < .05 (two-tailed). Otherwise, we used repeated measures ANOVA followed by Bonferroni-corrected pairwise treatment comparisons. For lickometer-generated data, custom scripts in Python were used to locate timestamps of lick bout onsets and offsets from Med Associates output data. Lick bouts were defined as a sequence of rapid licks (minimum of 3 licks with no more than 1000ms between each lick). Time-dependent data (latencies to commence lick bout and bout duration) were log-transformed for inferential statistics. For choice data, log-normal distributions were fit to empirical alcohol and grain RT density distributions. Response times (RTs) were log transformed for inferential statistics. We calculated two metrics from sampling RTs, the RT ratio and the winning RT. The RT ratio was calculated by dividing the log mean alcohol RT by the sum of the log mean grain and alcohol RT. The winning RT was generated by computing the proportion of pairwise alcohol-grain comparisons where the alcohol RT is faster than the grain RT. We fit simple linear regressions between these metrics during sampling trials and decision outcome in choice trials. Simple linear regressions were also fit between the mean and standard deviation of RT. A multiple regression was used to predict percentage alcohol choice in discrete choice from percentage alcohol choice in the home cage and during free operant selection. Simple linear regressions were performed for graphing purposes. All comparisons were paired sample t-tests (two-tailed) except when testing the effects of GLP1-ELP-FGF21, where an independent sample t-test (one-tailed) was used.

### Fiber photometry

Mice with histologically verified virus and cannula placement in the AcbSh were included for analyses (n = 7; Figure 4). Lick-related timestamps were extracted into a custom MATLAB pipeline alongside raw fibre photometry signals (410 nm isosbestic, 470 nm GRAB-DA3 DA-dependent). Fibre photometry signals were down-sampled (16.667 Hz), and invalid signals (e.g., movement artefacts, major non-event related transients outside the event-recording period) were excluded from further analysis. Remaining signals were low-pass (3 Hz) and band-stop (1.032-1.033 Hz, 2.547-2.551 Hz) filtered. The isosbestic signal was fitted onto the DA-dependent signal using robust regression (Keevers & Jean-Richard-Dit-Bressel, 2025) and dF/F signals calculated as: (470 nm signal – fitted 410 nm signal) / fitted 405 nm signal. Median detrend (60s window) was applied to the dF/F to remove non-phasic signal changes. dF/F peri-event signals were then aggregated, where 1.5 seconds before first lick and 3 seconds before bout onset and bout offset were used as a baseline, and 7 seconds following each event was analysed as the event-related transient.

Effects of treatment (GLP1-ELP-FGF21 agonist versus saline, exendin-4 versus saline) on event transients were analysed through cross-session event comparison of respective trials, where a difference transient was calculated. For both baselined and treatment comparison transients, a bootstrapping confidence interval (bCI; 95% CI, 1,000 boots) procedure was used to determine significant event-related activity, (Jean-Richard-dit-Bressel et al., 2020). Random resampling of trial signals with replacement, for the same number of trials, was used to construct a distribution of bootstrapped means. For each time-point the 2.5^th^ and 97.5^th^ percentiles of this distribution were used to determine a confidence interval, that was expanded by √(𝑛/𝑛−1) to account for narrowness bias. Periods of activity where the 95% CI did not include 0% dF/F for at least 0.33 s were considered significant.

### Evidence accumulation modelling

The Linear Ballistic Accumulator (LBA) (Brown & Heathcote, 2008) and Racing Diffusion Model (RDM) (Tillman et al., 2020) decompose RT distributions and choice probabilities into latent cognitive parameters which are informative about underlying choice processes. These latent cognitive parameters are accumulation rate for each response option (*v*), response caution (*B;* impulsiveness) and non-decision time (*t0:* motor and encoding time). In brief, these models propose a race between two competing choices with distinct accumulators that begin at some starting point (*A*). The first accumulator to reach a threshold (*b*) determines the choice made. These models primarily differ in their account of ‘noise’ in this decision-making process, where the LBA allows for between-trial variability in accumulation (*sv*) and the RDM allows for within-trial variability in accumulation (*s*). In the EMC2 package used to fit these data (Stevenson et al., 2024), models are therefore dependent on four shared parameters (*v*, *B*, *A*, *t0*) and a distinct variability parameter (*s* or *sv*). In EMC2, these parameters are estimated using Hierarchical Bayesian procedures where group-level parameters are fit using Gibbs sampling (Geman & Geman, 1984) and individual-level parameters using a particle Metropolis sampling algorithm (Gunawan et al., 2020). We used package default settings and priors with three independent Markov Chain Monte Carlo (MCMC) chains. Convergence was assessed using the Gelman-Rubin scale reduction factor (R^ < 1.1; Gelman and Rubin, 1992) and visual inspection of trace plots. The effective sample size (Eff > 100) was also assessed. Models were compared using the Deviance Information Criterion (DIC) (Spiegelhalter et al., 2002) and the Bayesian Predictive Information Criterion (BPIC) (Ando, 2011). Model fits were assessed using a Posterior Predictive check for the 97.5^th^, 50^th^ and 2.5^th^ percentile of RTs and for overall choice probability with 95% credible intervals (Heathcote et al., 2019). To make inferences about parameter estimates we generated a posterior probability value (Posterior p value). This was computed by taking the posterior distribution of the difference between treatments and estimating the posterior probability that this difference exceeded zero. Posterior p values < 0.025 were considered significant.

## Results

### Experiment 1. Alcohol consumption

Male mice were first maintained on a 4 hour intermittent alcohol access schedule, followed by four consecutive days of 4 hour daily access to alcohol to establish baseline intake prior to testing. Alcohol intake during this four-day baseline period did not differ between groups [mean ± SEM g/kg of 4 hour alcohol intake over the 4 days prior to test was 1.28±0.27 g/kg (Vehicle) and 1.37±0.32g/kg (GLP1-ELP-FGF21; F<1)].

On the test day, mice received an intraperitoneal (i.p.) injection of Vehicle or GLP1-ELP-FGF21 (1000 nmol/kg) and were subsequently given three sessions of overnight (24 hour) two-bottle choice alcohol access on an intermittent schedule (**Figure 1A**). GLP1-ELP-FGF21 significantly reduced overnight alcohol intake at both 24 hours (*F*(1,7) = 7.059, *p* = 0.033) and 72 hours post-injection (*F*(1,7) = 13.561, *p* = 0.008), with intake returning to baseline by 120 hours (*F* < 1; **Figure 1B**).

**Figure 1.**
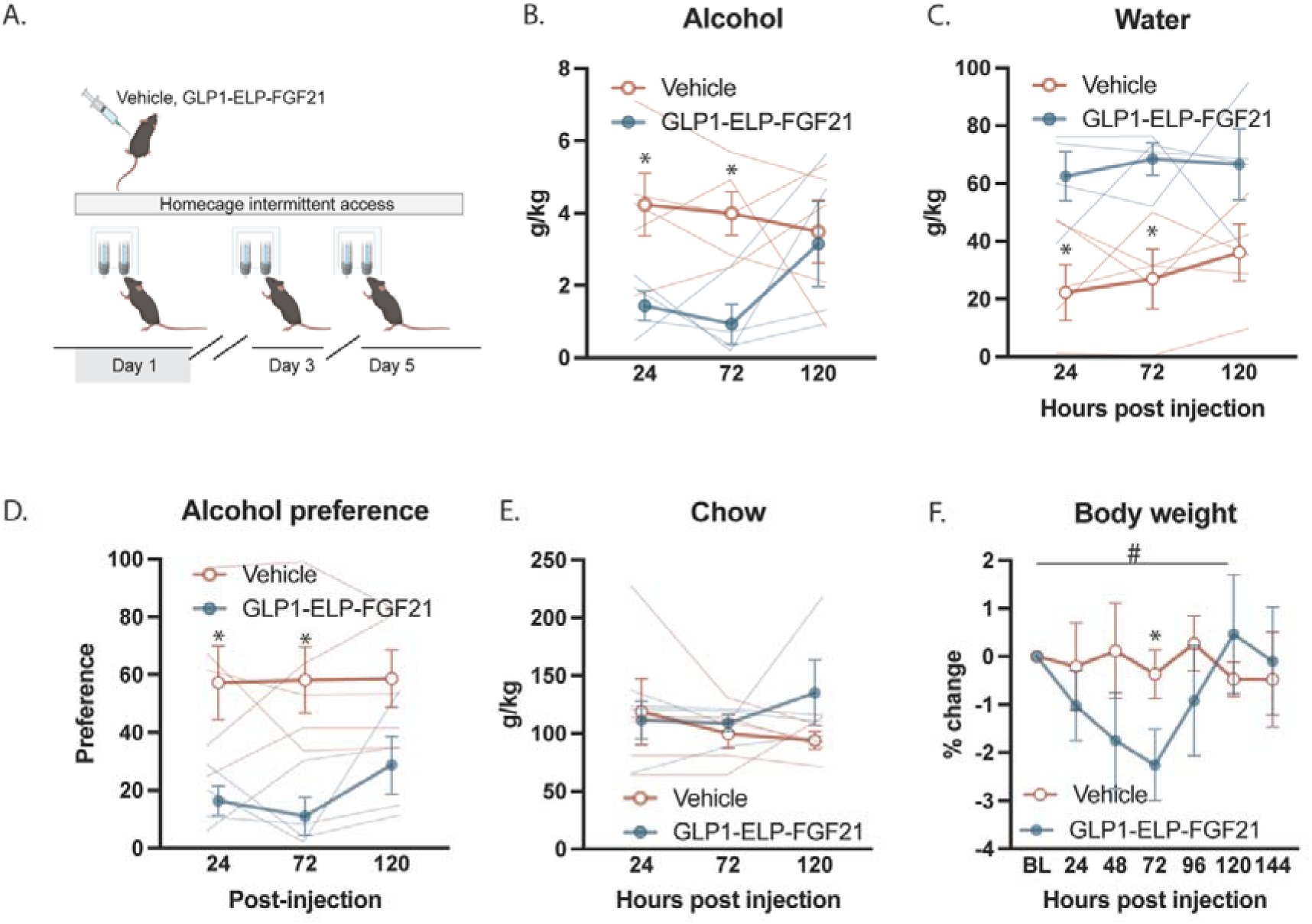
Homecage alcohol consumption. Schematic of test phase (A); male mice received two bottle choice to alcohol and water in the homecage on days 1 (injection day), 3, and 5 of the test phase. Effect of GLP1-ELP-FGF21 (1000nmol/kg) on overnight alcohol consumption (B), water consumption (C), alcohol preference (D), chow consumption (E), and bodyweight (change relative to the day prior to test) (F). Data are means±SEM. Figures 1A-E: \**p* < 0.05. Figure 1F: \**p* < 0.05 quadratic trend for GLP1-ELP-FGF21; #*p* < 0.05 group x quadratic trend analysis. *N* = 9 (Vehicle = 5; GLP1-ELP-FGF21 = 4).

### Water consumption

Conversely, GLP1-ELP-FGF21 treatment produced a significant increase in water intake. Water consumption (g/kg) was significantly elevated at 24 hours (*F*(1,7) = 7.806, *p* = 0.027) and 72 hours (intake: *F*(1,7) = 15.539, *p* = 0.006), but not at 120 hours post-injection day (*F*(1,7) = 4.853; *p* = 0.055; **Figure 1C**).

### Alcohol preference

Consistent with reduced alcohol intake, GLP1-ELP-FGF21 treatment significantly reduced alcohol preference at 24 hours (*F*(1,7) = 7.352, *p* = 0.03) and 72-hours (*F*(1,7) = 11.098, *p* = 0.013), with no significant effect at 120 hours post-injection (*p* = 0.075; **Figure 1D**).

### Food (chow) consumption

Overnight chow consumption (g/kg) was unaffected by GLP1-ELP-FGF21 treatment. Neither overall intake nor intake at individual post-injection time points (24, 72, or 120□hours) differed between groups (24, 72, and 120 hours: *p* values > 0.156; **Figure 1E**), indicating that reductions in alcohol intake were not attributable to nonspecific suppression of feeding.

### Body weight

GLP1-ELP-FGF21 treatment induced transient reduction in body weight, which was significant at 72 hrs following injection with (F(1,7) = 5.675, p = 0.049). Across the 7 day observation period, treated mice exhibited a pronounced U-shaped trajectory, characterized by initial weight loss followed by a recovery to baseline (quadratic trend contrast: F(1,7) = 10.219, p = 0.015). Vehicle-treated mice maintained stable body weight across this period (quadratic and linear trend contrasts: F values<1). The pattern of body weight changes differed significantly between groups (group x quadratic trend, F(1,7) = 8.042, p = 0.025; **Figure 1F**).

Together, these findings demonstrate that a single administration of GLP1-ELP-FGF21 results in a sustained reduction in alcohol intake and preference lasting up to 72 hours, accompanied by increased water consumption and transient weight loss, without affecting food intake.

### Experiment 2. Alcohol consumption during home cage intermittent access

Experiment 2 assessed the effects of GLP1-ELP-FGF21 and exenatide (a GLP-1 agonist), on AcbSh dopamine activity around alcohol intake in male mice (*n* = 11) maintained on an overnight intermittent access alcohol-schedule. Prior to drug testing, mice consumed an average of 3.55 g/kg/24 h of alcohol per mouse. Following administration of each treatment, mice underwent three intermittent sessions of overnight alcohol-access across 7 days, and alcohol intake was measured at 30, 100, and 148 hours post-injection of drug. There were significant main effects of treatment (*F*(2,10) = 5.233, p = 0.015), day (*F*(2,10) = 31.353, p<0.001) and a treatment x day interaction (*F*(4,10) = 5.723, p<0.01) on alcohol intake. Pairwise comparisons showed that GLP1-ELP-FGF21 reduced alcohol intake relative to vehicle at 30 hours (adjusted *p* = 0.049) and 100 hours (adjusted *p* = 0.02), and relative to exenatide at 100 hours (adjusted *p* = 0.042), indicating a sustained and reversible suppression of alcohol intake by GLP1-ELP-FGF21 treatment (**Figure 2A**). All other pairwise comparisons were not significant.

**Figure 2.**
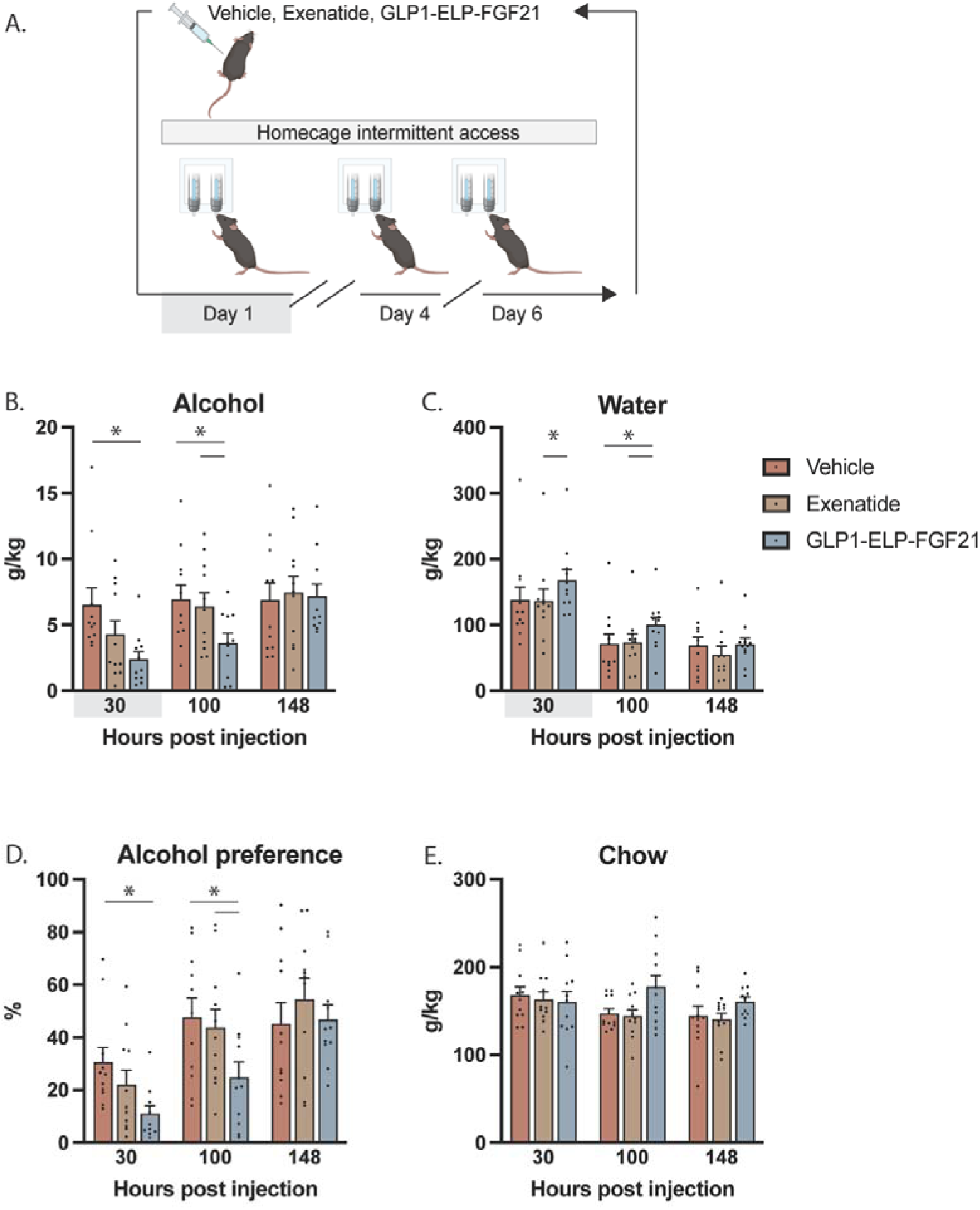
Homecage alcohol consumption. Schematic of test phase (A); male mice received two bottle choice to alcohol and water on days 1 (injection day), 4, and 6 during the test phase. Effect of exenatide (2.5μg/kg) and GLP1-ELP-FGF21 (1000nmol/kg) on alcohol consumption (B), water consumption (C), and alcohol preference (D) on homecage alcohol consumption test. Data are means±SEM. *p<0.05 (pairwise comparisons). *N* = 11.

### Water consumption

Water intake showed significant main effects of treatment (*F*(2,10) = 8.925, p =.002), day (*F*(2,10) = 71.722, *p* < .001), with no significant treatment x day interaction (*F*(4,10) = 2.383, *p* = 0.068). Pairwise comparisons showed that GLP1-ELP-FGF21 treatment increased water consumption relative to vehicle at 100 hours (adjusted *p* = 0.010) and relative to exenatide at both 30 hours and 100 hours (adjusted *p* = 0.014 and 0.001, respectively; **Figure 2C**). All other pairwise comparisons were not significant.

### Alcohol preference

There was an overall main effect of treatment on alcohol preference (*F*(2,10) = 6.816, *p* = 0.006), day (*F*(2,10) = 39.305, *p* < 0.001) and treatment x day interaction (*F*(4,10) = 4.565, *p* = 0.004). Pairwise comparisons showed reduced alcohol preference following GLP1-ELP-FGF21 treatment relative to vehicle at 30 hours and 100 hours (adjusted *p* = 0.048 and 0.010, respectively), and relative to exenatide at 100 hours (adjusted *p* = 0.016; **Figure 2D**). All other pairwise comparisons were not significant.

### Chow intake

There was a main effect of treatment on chow intake (*F*(2,10) = 4.078, *p* =.033), no main effect of day (*F*(2,10) = 1.831, *p* =.186) and a significant treatment x day interaction (*F*(4,10) = 2.91, p=.033). Pairwise comparisons revealed increased chow intake following GLP1-ELP-FGF21 relative to exenatide at 100 hours (adjusted *p* = 0.007; **Figure 2E**). All other pairwise comparisons were not significant.

### Body weight

Across all treatments body weight increased significantly from post-injection day 1 to day 6 (linear trend: *F*(1,10) = 21.655, *p* = 0.001; relative to body weight on the day of drug treatment), consistent with recovery from water deprivation on the test day. There were no significant differences between treatment groups (main effect, treatment x day; pairwise comparisons; all F < 1.447, *p* > 0.257).

### Lick microstructure and fiber photometry recording of accumbal dopamine activity

To examine how GLP1-ELP-FGF21 modulates dopamine signaling during alcohol consumption, a subset of mice (*n* = 7) underwent fiber-photometry recording of nucleus accumbens dopamine activity during single-bottle lickometer access to alcohol prior to their overnight home-cage alcohol-access sessions (Gibson et al., 2018).

Analysis of lick microstructure revealed significant treatment effects on bout size quantified by number of licks per bout (*F*(2,6) = 3.958, *p* = 0.048, **Figure 3A**) and latency to initiate the first bout (*F*(2,6) = 9.698, *p* = 0.003, **Figure 3F**). Pairwise comparisons of treatments showed significant differences in latency to initiate the first bout between GLP1-ELP-FGF21 and exenatide (adjusted *p* = 0.011), and between exenatide and vehicle (adjusted *p* = 0.022; **Figure 3F**). All other pairwise comparisons were not significant (adjusted *p* > 0.098).

**Figure 3.**
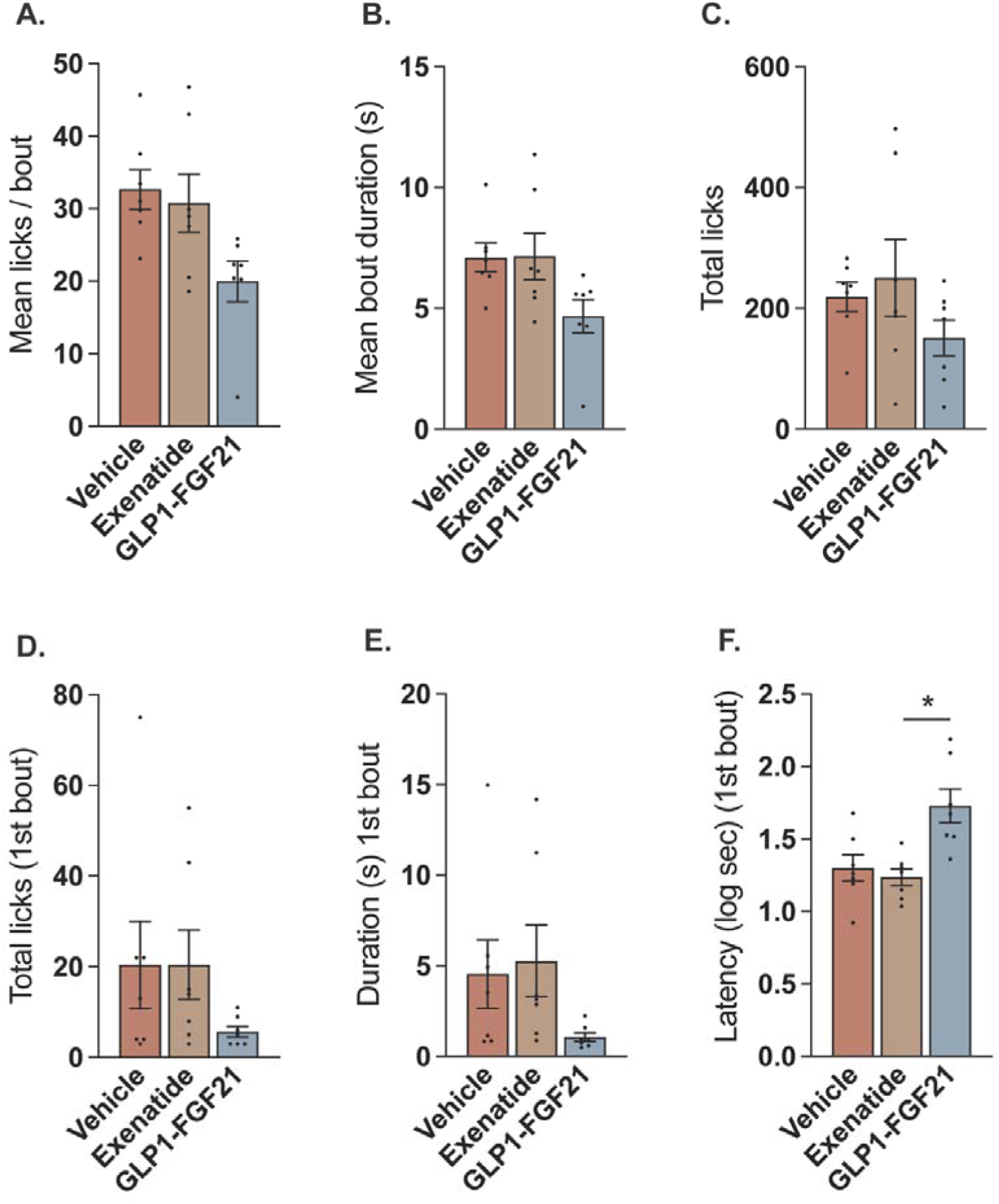
Lick microstructure. Effect of exenatide (2.5μg/kg) and GLP1-ELP-FGF21 (1000nmol/kg) on lick frequency per bout (A), bout duration (B), total licks (C), lick frequency during first lick bout (D), duration of first lick bout (E), and log-transformed latency to initiate first lick bout (F) in male mice during brief (20 min) lickometer access to alcohol on test. Data are means±SEM. *p<0.05. *N* = 11.

Fibre photometry recording using GRAB-DA3 revealed robust modulation of dopamine activity at lick-bout onset and offset under vehicle treatment. **Figure 4** shows the mean ±□95% bootstrapped confidence intervals. Under vehicle treatment, DA activity was robustly modulated following the starts (lick bout onset; **Figure 4E**) and ends (lick bout offset; **Figure 4F**) of alcohol drinking. In contrast, under treatment with either GLP1-ELP-FGF21 or exenatide, this modulation was either diminished or abolished (**Figures 4E-F**). Within-subject treatment comparison indicated significant differences in DA transient activity between vehicle and GLP1-ELP-FGF21 in the seconds following the onset and offset of a lick bout (**Figures 4E-F**). Under exenatide, similar comparison also revealed significantly different DA transient activity from vehicle however these periods were delayed and less robust (**Figures 4E-F**). On examining DA transient around first lick, isolated from the influence of prior ingestion of alcohol, DA transient was significantly modulated under both vehicle and GLP1-ELP-FGF21, however these modulations were significantly distinct from one another (**Figure 4G**). Transient DA activity around first lick was excitatory following vehicle treatment. In contrast, following treatment with GLP1-ELP-FGF21, DA activity was negatively modulated following the first lick.

**Figure 4.**
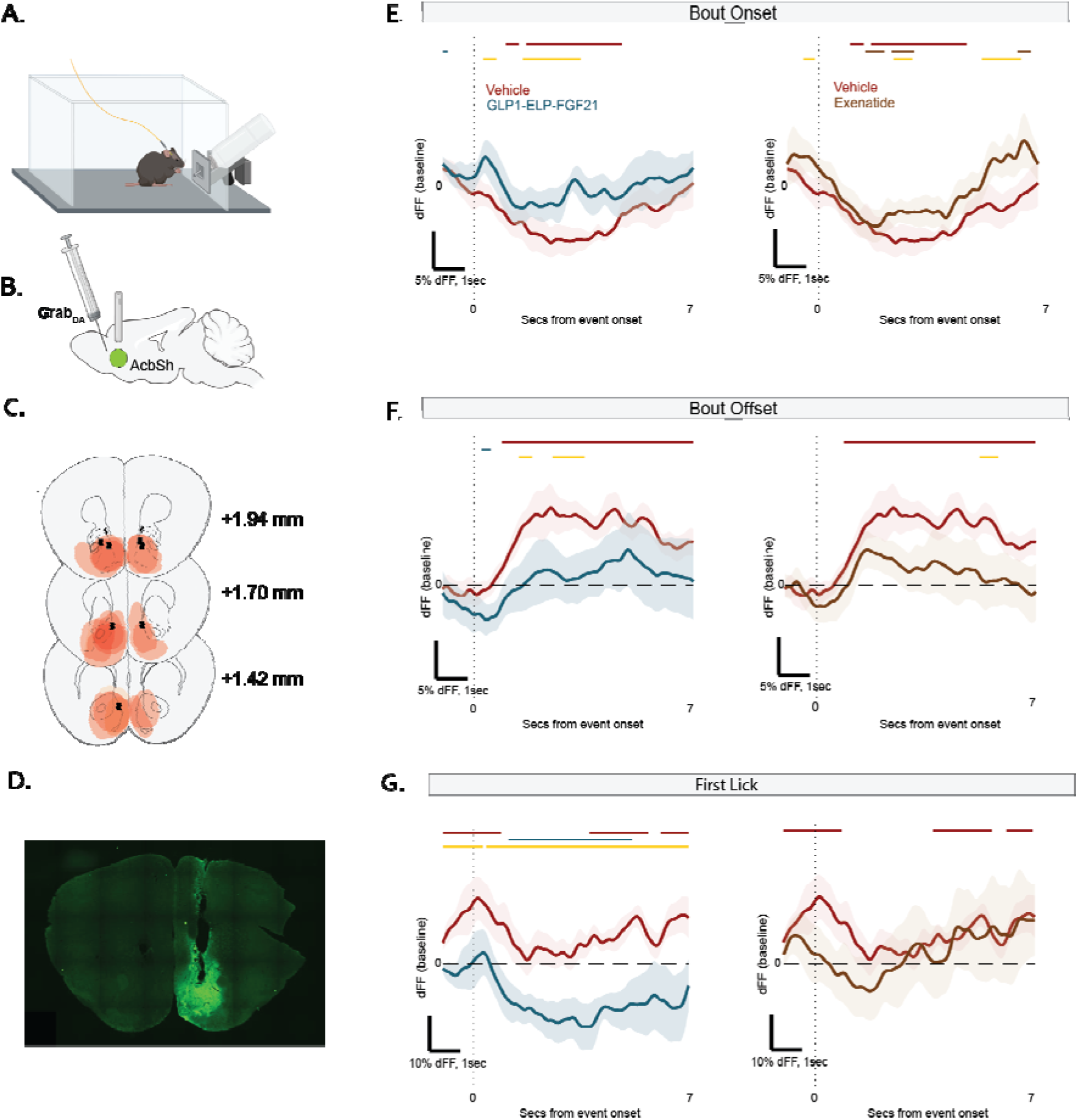
Accumbal DA activity. Fiber photometry recording of DA activity in nucleus accumbens. Schematic of lickometer task (A). Unilateral intracranial delivery of GRAB-DA3 sensor and optic cannula in Acb (B). Histology-verified location of optic cannulae and viral spread; placements were counterbalanced across hemispheres (C). Histomicrograph of example cannulae position and viral infection (D); Effect of exenatide (2.5µg/kg) and GLP1-ELP-FGF21 (1000nmol/kg) on DA activity modulated around bout onset (E), bout offset (F), and first lick (G) in male mice during brief (20 min) lickometer access to alcohol on test. Data are means±SEM. *p<0.05. *N* = 7.

### Experiment 3. Effect of GLP1-ELP-FGF21 in female mice

Experiment 3 evaluated the effects of GLP1-ELP-FGF21 in female mice (*n* = 10) across four doses: 0, 100, 300, and 1000 nmol/kg. One mouse was excluded from water intake analyses due to a water bottle leak; therefore, water consumption and alcohol preference analyses were conducted with n = 9.

### Alcohol Consumption

A repeated-measures ANOVA revealed a significant effect of GLP1-ELP-FGF21 dose on alcohol intake (*F* (3,27) = 4.415, *p* = 0.012; **Figure 5A**). Pairwise comparisons showed that the highest dose of 1000 nmol/kg significantly reduced alcohol consumption at 24 hours post-injection compared with all other doses (adjusted *p* < 0.029; **Figure 5A**). No other pair-wise comparisons were significant (*p* > 0.259).

**Figure 5.**
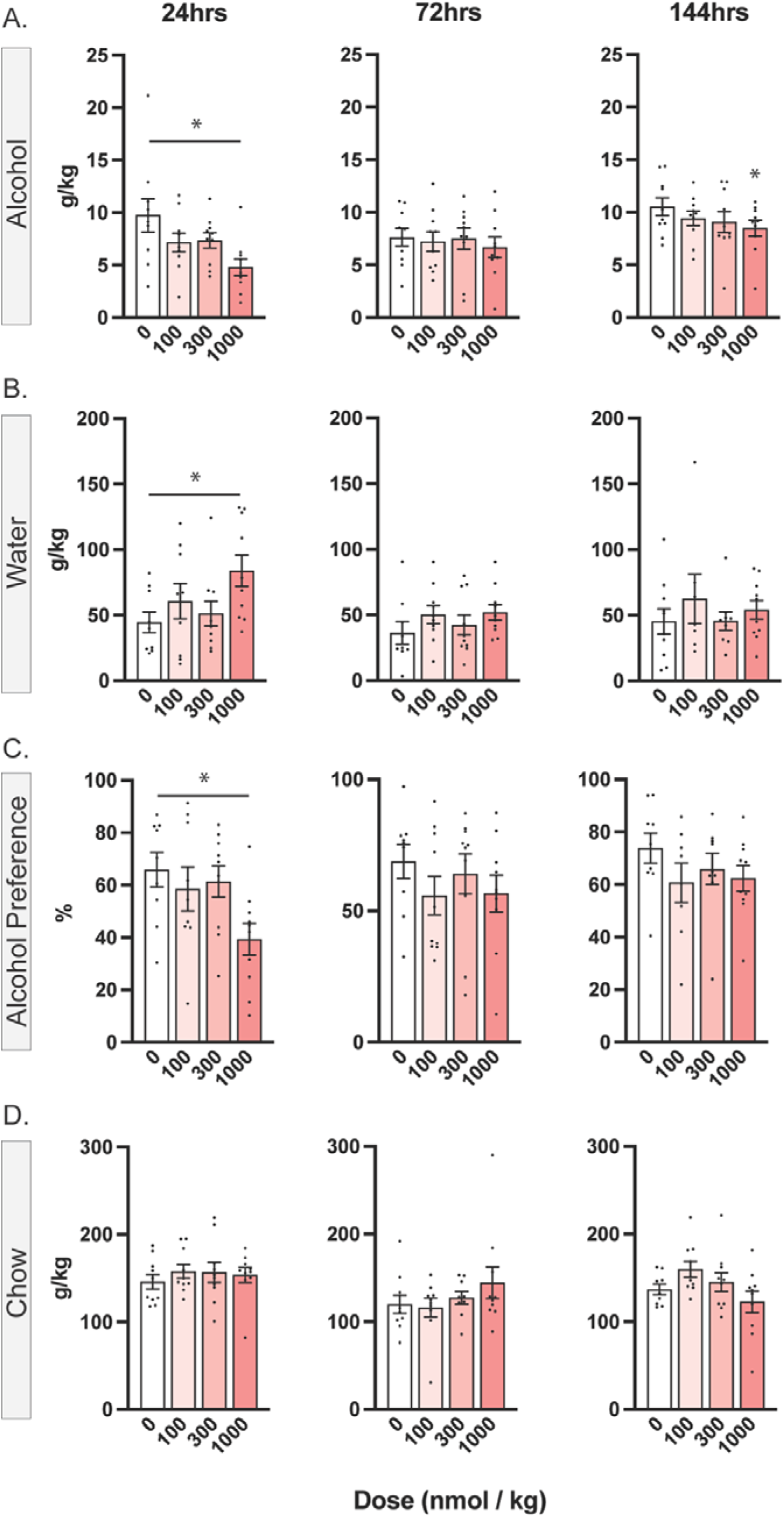
Homecage alcohol consumption (females). Effect of GLP1-ELP-FGF21 (1000nmol/kg) in female mice on alcohol consumption (A), water consumption (B), alcohol preference (C), and chow consumption (D), across 24 hours (left), 72 hours (centre) and 144 hours (right) post-injection. Data are means±SEM. \**p* < 0.05 pairwise comparisons relative to vehicle. *N* = 9-10.

### Water Consumption

At 24 hours post-injection, the highest dose (1000 nmol/kg) significantly increased water consumption relative to vehicle (adjusted *p* = 0.025; **Figure 5B**). No other dose comparisons reached significance (*p* > 0.073).

### Alcohol Preference

Pairwise comparisons showed that the 1000 nmol/kg dose significantly reduced alcohol preference at 24 hours post-injection of GLP1-ELP-FGF21 relative to vehicle (adjusted *p* = 0.025) and the 300 nmol/kg dose (adjusted *p* = .018; **Figure 5C**). All other effects on alcohol preference were not significant (*F* < 3.065).

### Chow Consumption

GLP1-ELP-FGF21 did not alter chow intake at any dose across 24, 72, or 144 hour time points (*p* > 0.122; **Figure 5D**).

### Bodyweight

No significant effects of GLP1-ELP-FGF21 on body weight were observed at any dose or time point (*p* > .856).

### Experiment 4. Effect of GLP1-ELP-FGF21 on Alcohol Choice

Male mice received intermittent home-cage alcohol access prior to self-administration training. Mean ±SEM intake levels for overnight and 2 hour access shown in **Table 1**.

**Table 1:** Home cage consumption for Experiment 4 in g/kg.

|  | 24 hr intermittent access | 2-hr intermittent access |
| --- | --- | --- |
| All non-drug sessions | $8.14 \pm 0.54$ | $0.73 \pm 0.06$ |
| GLP1-ELP-FGF21 agonist | $8.26 \pm 0.66$ | $0.69 \pm 0.07$ |
| Devaluation | $5.15 \pm 0.90$ | $0.70 \pm 0.12$ |

### Discrete-choice behavior aligns with evidence-accumulation frameworks

We first assessed discrete-choice behaviour in the absence of pharmacological testing (n = 20, males; **Table 2**). Mice displayed a significant overall preference for grain over alcohol (t(1,19) = 3.669, *p* < 0.01, **Figure 6B**) with no difference in log-mean reaction times (RTs) between options (p = 0.76, **Figure 6C**). RTs were rapid for both choices (Alcohol median = 1.6 seconds, Grain median = 1.2 seconds) and followed positively skewed log-normal distributions (Alcohol R^2^ = 0.97, F(1,98) = 2866, *p* < 0.01; Grain: R^2^ = 0.98, F(1,98) = 4514, p< 0.01, **Figure 6D**). Linear coefficients of variation further supported log-normality (Alcohol: R^2^ = 0.90, F(1,18) = 169.3, *p* < 0.01; grain: R^2^ = 0.95, F(1,19) = 366, *p* < 0.01, **Figure 6D inset**).

**Figure 6:**
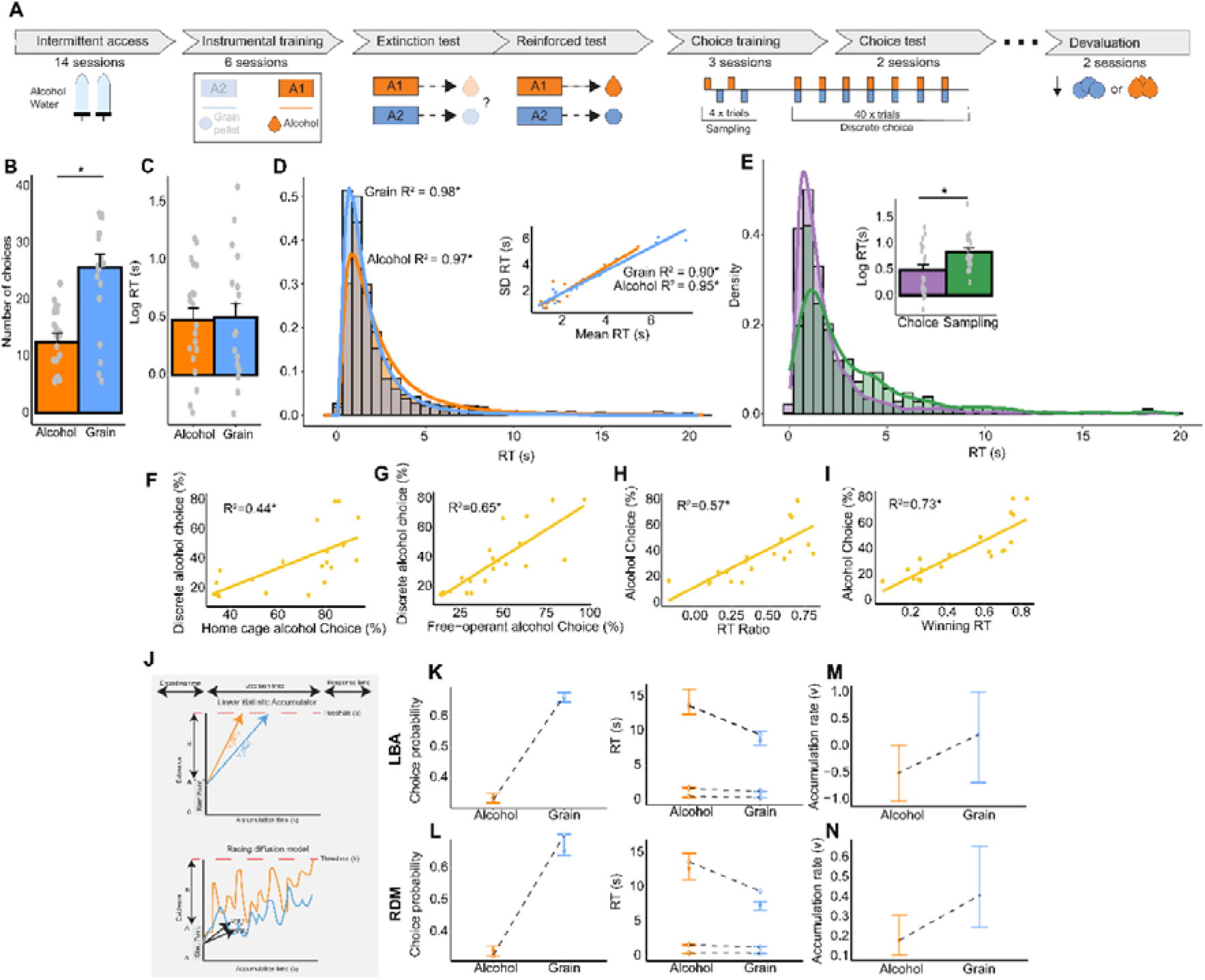
Discrete choice behaviour. Discrete choice alcohol behaviour and cognitive modelling. Schematic of behavioural procedures (A). Instrumental choice (B) and log RTs (C) for all non-drug choice sessions. Fits of RTs to log-normal distributions and linear regression of mean RTs and standard deviations (D). Distributions and log RTs of sampling and choice (E). Simple linear regressions of discrete alcohol preference (%) with home cage preference (%) (F) and free-operant preference (%) (G). Simple linear regressions of discrete alcohol preference (%) with RT ratio (H) and Winning RT (I). Schematic of the LBA and RDM framework (J). Posterior predictive fits of choice probabilities and RTs for LBA (K) and RDM output (L). Open and closed circles represent observed and predicted data. Estimated accumulation rates for alcohol and grain choice in the LBA (M) and RDM (N). Data are means±SEM, except for accumulation rates where they are posterior medians with 95% credible intervals. *p<0.05 or *Posterior p< 0.025. *N* = 20.

**Table 2:** GLP1-ELP-FGF21 on alcohol choice.

| Experiment | N | Cohort 1 - History | Cohort 2 - History |
| --- | --- | --- | --- |
| All non-drug sessions | 20 | Behavioural training, Drug testing | Behavioural training, Drug testing, Devaluation |
| GLP1-ELP-FGF21 treatment | Vehicle = 6, GLP1-ELP-FGF21 = 8 | Behavioural training | Behavioural training |
| Devaluation | 8 | N/A | GLP1-ELP-FGF21 treatment |

Alcohol choice in the discrete choice task was strongly predicted by home-cage alcohol preference and free-operant alcohol choice (R^2^_adj_ = 0.72, F(3,16) = 17.63, p < 0.01, **Figure 6F-G**). There were no interaction between predictors (*p* = 0.059).

### Behavioural signatures of evidence accumulation

We next tested whether discrete choice behaviour conformed to predictions of evidence accumulation models. Two predictions were confirmed. First, RTs during choice trial were shorter than during sampling trials (sequential encounters) [t(1,19) = 4.61, p < 0.01, **Figure 6E**] consistent with cross-censoring of latency distributions (Vandaele et al., 2021). Second, sampling trials RT predicts later choice. Two metrics generated from sampling trials RTs—the RT ratio (R^2^ = 0.57, p < 0.01, **Figure 6H**) and the winning RT (R^2^ = 0.73, p < 0.01, **Figure 6I**) strongly predicted subsequent choice. This suggests a theoretical grounding for applying formal evidence accumulation models to this task.

To formalise these behaviours, we modelled the relationship between choice probability and the distribution of RTs using two evidence accumulation race models: the Linear Ballistic Accumulator (LBA) and Racing Diffusion Model (RDM) (**Figure 6J**). In both models we allowed the latent cognitive parameter, accumulation rate (*v*) to vary with choice and estimated non decision time (*T0*), starting point (*A*), caution (*B*) across choices. Information criteria, a goodness-of-fit metric only slight favoured the RDM variant (DIC: LBA = 26298, RDM = 26230; BPIC: LBA = 26458, RDM = 26403) and thus we report the results of both model parameterisations.

Both models fit the data well as indicated by the overlap of predicted credible intervals and observed choice probabilities, and RTs with minor overestimation at longer RTs in the RDM (**Figure 6K-L**). Models revealed marginally slower accumulation rates for alcohol choices relative to grain (LBA: Posterior p = 0.058; RDM: Posterior p = 0.027 **Figure 6M-N**).

### GLP1-ELP-FGF21 slows the accumulation rate of alcohol rewards during choice

We next tested the effects of GLP1-ELP-FGF21 on reward choice (Vehicle: n = 6, GLP1-ELP-FGF21: n = 8). On the final day of discrete choice training, alcohol preference did not differ between counterbalanced treatment groups (p = 0.43). When tested under GLP1-ELP-FGF21 treatment, mice made fewer alcohol choices (t(1,12) = 2.02, p = 0.033, **Figure 7A inset**) without altering grain choices (p = 0.707, **Figure 7B inset**). Although RTs between alcohol or grain choices did not differ between treatments (p > 0.342), the RT distribution showed a downward shift in its peak, suggesting a potential change in evidence accumulation dynamics under treatment (**Figure 7A**).

**Figure 7:**
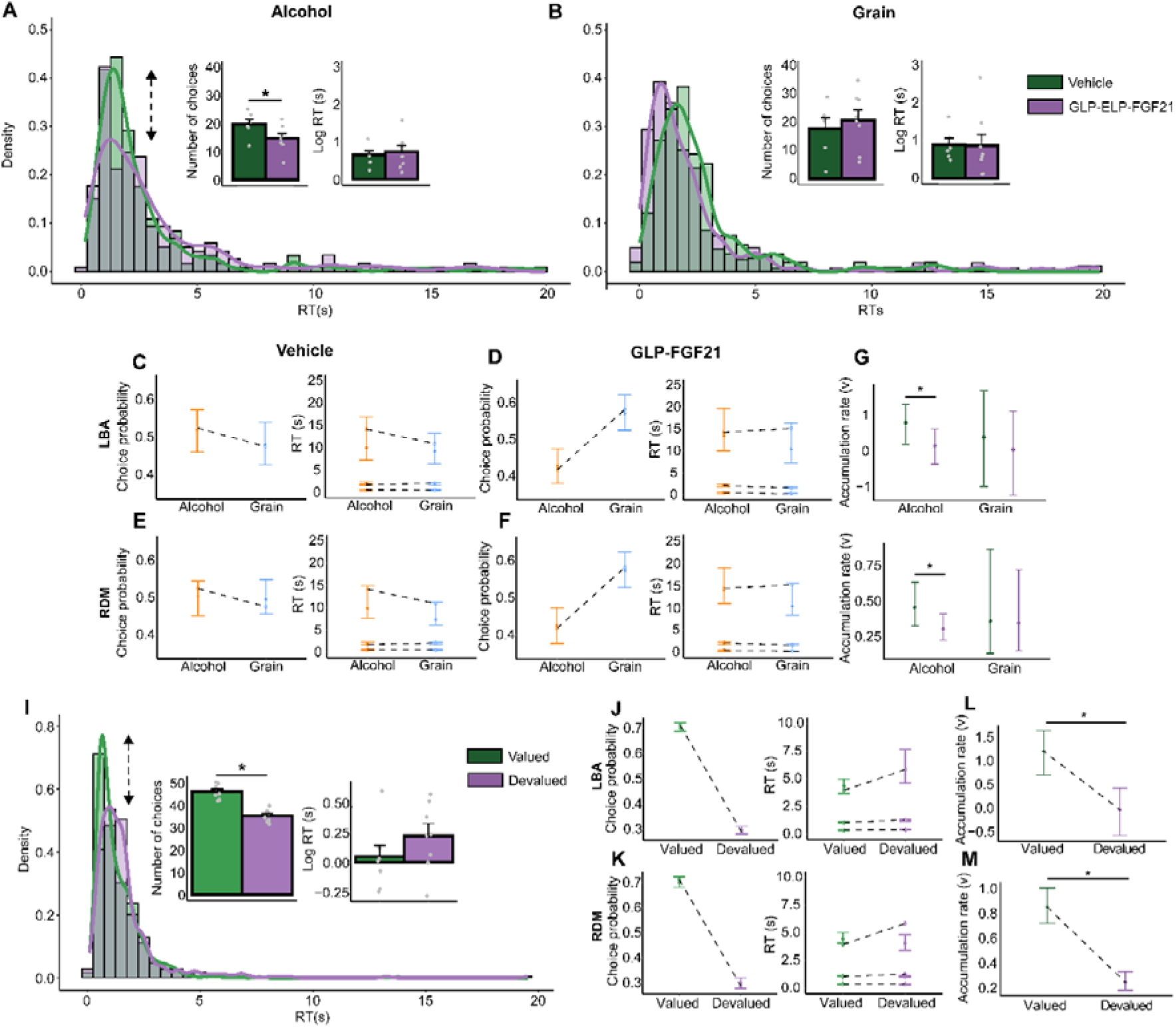
GLP1-ELP-FGF21 on alcohol choice. Effect of GLP1-ELP-FGF21 (1000nmol/kg) and sensory-specific satiety on discrete choice alcohol behaviour. Instrumental responding, log RTs and distributions under vehicle and GLP-ELP-FGF21 for alcohol (A) and grain (B) choices. Posterior predictive fits of choice probabilities and RTs under vehicle (C) and GLP-ELP-FGF21 (D) for the LBA. Posterior predictive fits of choice probabilities and RTs under vehicle (E) and GLP-ELP-FGF21 (F) for the RDM. Open and closed circles represent observed and predicted data. Estimated accumulation rates of alcohol and grain choices for the LBA (G) and RDM (H). Instrumental responding, log RTs and distributions valued and devalued responses (I). Posterior predictive fits of choice probabilities and RTs for the LBA (J) and RDM (K). Estimated accumulation rates for valued and devalued responses for the LBA (L) and RDM (M). Data are means±SEM, except for accumulation rates where they are posterior medians with 95% credible intervals. *p<0.05 or *Posterior p two tailed< 0.025, one tailed< 0.05. Pharmacological treatment, GLP-ELP-FGF21 = 8, Vehicle = 6. Sensory specific-satiety, N = 8.

To formally decompose the joint choice-RT distribution into latent cognitive variables, data for the treatment and vehicle groups were separately fit to the LBA and RDM models. Information criteria showed no strong difference between models in either condition (DIC: LBA = 2263, RDM = 2268; BPIC: LBA = 2386, RDM = 2402) and treatment models (DIC: LBA = 2853, RDM = 2818; BPIC: LBA = 2997, RDM = 2928). Both models fit the data well as indicated by the close overlap between predicted credible intervals and observed choice probabilities and RTs (**Figure 7C-F**).

Importantly, following GLP1-ELP-FGF21 treatment, there was reliably slower accumulation rates for alcohol (LBA: Posterior p = 0.043, RDM: Posterior p = 0.035) but not grain choices (LBA: Posterior p = 0.34 RDM: Posterior p = 0.48 **Figure 7G-H**). There were no differences in any other model parameters (all comparisons: LBA, Posterior p > 0.25, RDM, Posterior p > 0.20; data not shown). Together, this suggests a specific effect of GLP1-ELP-FGF21 on decision making architecture by attenuating the accumulation rate for alcohol choices.

### Changes in value are driven by accumulation rate during choice

Finally, while it has been suggested that changes in reward value are related to accumulation rate in human value-based decision tasks (Busemeyer & Townsend, 1993; Clithero, 2018) this has not been shown empirically in animals. Thus, to confirm that changes in reward value are recapitulated by accumulation rates, we took a subset of animals (n = 8) and devalued rewards using within-subjects manipulations of sensory specific satiety. This was successful as animals chose the devalued reward less than the valued reward (t(1,7) = 5.21, p < 0.01, **Figure 7I inset**). While there was no difference in the RTs between valued and devalued rewards (p = 0.172) we note the lower peak in the RT distribution of the devalued reward suggesting changes in accumulation rate (**Figure 7I**).

We modelled the joint relationship between choices for devalued and valued rewards and distribution of RTs using the RDM and LBA with model parametrizations as previously noted. Information criteria only marginally favoured the LBA variant (DIC: LBA = 6482, RDM = 6534; BPIC: LBA = 6629, RDM = 6676) and thus we report the results of both models. Both fit the data well as indicated by the overlap of predicted credible intervals and observed choice probabilities and RTs (**Figure 7J-K**) but we note some misfits in the RDM model. Interestingly, there were reliably slower accumulation rates for devalued choices (LBA: Posterior *p* < 0.01; RDM: Posterior *p* <0.01 **Figure 7L-M**) suggesting that manipulations of reward value are consistent with changes in evidence accumulation rate at the time of choice.

## Discussion

In this study, we present preclinical findings across behaviour, cognitive modelling and fiber photometry tracking of dopamine activity in alcohol drinking mice, supporting GLP1-ELP-FGF21 to attenuate alcohol consumption and highlighting its potential as a novel treatment strategy for AUD.

The established safety profile, clinical availability and widespread adoption of GLP-1 agonists provide an advantageous platform for its repurposing for AUD. Unimolecular multi-agonist hybrid approaches are now emerging and even approved for use (e.g. tirzepatide), leveraging complementary mechanisms to augment efficacy. However, much of this development has been confined to hybrid peptides within the glucagon superfamily (e.g. GLP-1, GIP, GCGR), exploiting sequence similarities and convergent signalling pathways (Clemmensen et al., 2019).

In contrast, GLP1-ELP-FGF21, enables dual targeting of two distinct yet complementary peptide pathways–GLP-1 and FGF21, via conjugation to an ELP linker. The intrinsically disordered, temperature-sensitive ELP domain provides sustained zero-order release of the drug for approximately one week (Gilroy et al., 2020). This sustained release profile arises from the temperature-triggered phase behavior imparted by the ELP to the ternary fusion protein. The ELP was engineered to exhibit a lower critical solution temperature (LCST) phase transition at 30 °C, ensuring that the GLP1-ELP-FGF21 fusion remains soluble at room temperature in a syringe but undergoes a soluble-to-insoluble LCST transition upon subcutaneous injection. Once injected, the fusion protein forms an insoluble coacervate depot under the skin.

Drug is released from this depot with zero-order kinetics over 5–7 days as the fusion protein gradually dilutes at the depot margins. Dilution reverses the phase transition because the LCST is inversely related to ELP concentration (Meyer & Chilkoti, 2004). This concentration-dependent phase behavior enables controlled, long-acting release of the therapeutic.

While the durable and cooperative properties of GLP1-ELP-FGF21 have been characterised in the diabetic-obese *Db/db* mouse model on measures of glycaemia and body weight (Gilroy et al., 2020), here, we provide the potential application of this dual agonist in relation to alcohol-motivated behaviour.

In mice with a history of chronic intermittent access to alcohol, GLP1-ELP-FGF21 significantly reduced alcohol consumption in male and female mice, consistent with mono-peptide agonism of GLP-1 and FGF21 in preclinical animal models of alcohol consumption and preference (Egecioglu et al., 2013; Flippo et al., 2022; Vallöf et al., 2016). While we did not test each peptide in isolation, prior work comparing GLP1-ELP-FGF21 against ELP-linked monotherapy (GLP1-ELP and FGF21-ELP) in *Db/db* mice indicated at least additive contributions of combining GLP1 and FGF21 (Gilroy et al., 2020), and possibly, similar additive contributions underlie the effects observed here in alcohol drinking mice.

Moreover, this reduction was long lasting following a single injection at the highest dose in male mice, though there was reduced potency when tested in females. Sex-dependent effects on alcohol intake have similarly been reported for GLP-1R agonists, for example dulaglutide had more pronounced and long-lasting effects on alcohol drinking in males than in females (Vallöf et al., 2020). In contrast, others have reported more pronounced effects on alcohol drinking in females versus in males following some doses of semaglutide (C. Aranäs et al., 2023).

While the mechanisms behind the observed sex difference in response to GLP1-ELP-FGF21 remain unclear, we also note experimental differences in our male and female mouse studies may have contributed, at least in part, to these differences, and further work is warranted to determine the relative contribution of sex to these effects.

In considering potential non-specific contributions to the reductions in alcohol intake, there was no reduction in home-cage food intake or lever pressing for grain pellets, indicating that the inhibitory effects of GLP1-ELP-FGF21 on alcohol drinking were unlikely driven by malaise – a common side effect of GLP-1R agonists, or sedation. Moreover, we did not see treatment-related reductions in body weight under conditions in which mice were water-restricted; reductions in weight gain were transiently observed in mice maintained under standard home cage conditions and the timing of these reductions were largely consistent with prior work involving GLP1-ELP-FGF21 (Gilroy et al., 2020). Indeed, reductions in bodyweight have been reported across studies involving GLP-1R agonists in alcohol drinking rats (C. Aranäs et al., 2023; Marty et al., 2020; Vallöf et al., 2020), but not well documented for FGF21 in alcohol drinking subjects, except in Flippo et al. (2022) where neither FGF21 nor its long-acting analogue reduced body weight in alcohol-drinking vervet monkeys.

The mechanisms through which GLP-1 and FGF21 regulate alcohol-motivated behaviour have been linked to signalling within the amygdala and mesolimbic dopamine system. Previous microdialysis studies have shown that acute systemic administration of alcohol increases extracellular accumbal dopamine release, and that this release was attenuated by GLP-1R agonists, semaglutide, liraglutide and exenatide (C. Aranäs et al., 2023; Egecioglu et al., 2013; Vallöf et al., 2016).

Here, we extend these findings, leveraging the spatiotemporal resolution afforded by the genetic engineered dopamine sensor to record dynamic modulations in dopamine release/binding while mice engaged in voluntary alcohol consumption. Following the initiation and termination of alcohol drinking, dopamine transients decreased and increased, respectively. Critically, these modulations were attenuated following treatment with the dual-agonist GLP1-ELP-FGF21, suggesting a decoupling in the relationship between dopamine and alcohol-consumption per se. That dopamine transients overall decreased following initiation of alcohol drinking is consistent with prior studies involving similar recordings of dopamine activity in the medial AcbSh around alcohol consumption and may be characteristic of medial AcbSh relative to adjacent accumbal subregions such as the accumbens core, where dopamine transients appear to instead increase around the consumption of alcohol (Liu et al., 2020). Whether and how GLP1-ELP-FGF21 alters specific signatures of dopamine transmission around alcohol consumption and other-alcohol related behaviours such as relapse remain to be determined. Nonetheless, we found that GLP1-ELP-FGF21 also altered, albeit modestly, the microstructure of alcohol licking, indicative of possible changes in the palatability and value of alcohol, and consistent with the role of GLP-1 receptors that regulate size and duration of lick bouts for nutritive palatable rewards such as sucrose (Dossat et al., 2013).

We also introduced a discrete choice alcohol task in mice. Consistent with broader literature we show that animals generally prefer the alternate reward over drug rewards and decide quickly between choice options (Augier et al., 2018; Cantin et al., 2010). Our results suggest that the dynamics of this decision-making are well-accounted for by evidence accumulation processes as they are consistent with general predictions of this framework (Freidin et al., 2009; Ojeda et al., 2018; Shapiro et al., 2008; Vandaele et al., 2021). Evidence accumulation models offer a mathematical account of decision-making with a generalisable solution across domains and species (Forstmann et al., 2016; Philiastides & Ratcliff, 2013) implying shared cognitive laws of decision-making (Eun et al., 2022). This is important given the converging evidence that addiction has been framed as a disorder of decision-making (Field et al., 2020; Hogarth, 2020; Lüscher et al., 2020) and the need to address mounting failures in the translation of pharmacological treatments to clinical populations (Heilig et al., 2019; Venniro et al., 2020).

We leverage these shared computations by reverse-translating two race models of evidence accumulation, the LBA and the RDM (Brown et al., 2017; Tillman et al., 2020). We find that these models provide a comprehensive and accurate account of the joint relationship between behavioural allocation and the characteristics of the RT distribution in discrete alcohol choice. They account well for canonical devaluation by sensory specific satiety via slowing in the latent accumulation rate of decision-making. In line with human data (Busemeyer & Townsend, 1993; Krajbich et al., 2012) our data suggest that evidence accumulation effectively operates in animal value-based tasks, likely sampling central value representations to evaluate courses of action (Busemeyer et al., 2019; Clithero, 2018).

We show that *GLP1-ELP-FGF21* reduces alcohol choice and shifts its temporal dynamics without influencing decision-making around grain. Formal modelling revealed that slowing in alcohol accumulation rate explained these observed behavioural effects well. Together, this suggests that *GLP-ELP-FGF21* likely drives a reduction in the computational value of alcohol at the time of choice without impacting the value of grain rewards. Such cognitive inferences are directly translatable to clinical research as analogous value-based decision-making tasks in clinical populations (Copeland et al., 2023; Hardy & Hogarth, 2017) can be leveraged using evidence accumulation models as a computational bridge (Redish et al., 2021).

The precise pharmacological determinants of our findings remain unclear. Both FGF21 and GLP-1 robustly modulate alcohol consumption and preference (Cajsa Aranäs et al., 2023; Egecioglu et al., 2013; Flippo et al., 2022; Schumann et al., 2016). GLP-1 also influences other alcohol-related behaviours such as conditioned place preference (Egecioglu et al., 2013), relapse-like drinking (Cajsa Aranäs et al., 2023; Chuong et al., 2023; Thomsen et al., 2017) and operant self-administration (Díaz-Megido & Thomsen, 2023; Egecioglu et al., 2013). The role of FGF21 in similar alcohol-motivated behaviours remain largely unexplored. As FGF21 and GLP-1 exhibit complementary and interdependent mechanisms of action in regulating weight-loss and insulin sensitivity (Gilroy et al., 2020; Liu et al., 2021; Liu et al., 2019; Pan et al., 2021; Yang et al., 2012) disentangling their specific contributions to alcohol motivated behaviours and alcohol choice will be important for future studies.

Taken together the present behavioural, cognitive and photometry experiments provide insights into the potential for GLP1-ELP-FGF21 on alcohol-related behaviour, tentatively via changes in underlying decision-making processes and dopamine mechanisms within the Acb. These data may provide a rationale for further investigations involving combined GLP-1 and FGF21 in treatment-seeking alcohol drinkers.

## Acknowledgements

Supported by a Synergy Grant from the National Health and Medical Research Council of Australia (GNT 2009851).

## Notes

### Competing Interest Statement

The authors have declared no competing interest.

### Summary of Updates

This version has been updated with corrections to choice modelling.

